# A comprehensive co-expression network analysis in *Vibrio cholerae*

**DOI:** 10.1101/2020.02.07.939611

**Authors:** Cory D. DuPai, Claus O. Wilke, Bryan W. Davies

## Abstract

Research into the evolution and pathogenesis of *Vibrio cholerae* has benefited greatly from the generation of high throughput sequencing data to drive molecular analyses. The steady accumulation of these datasets now provides a unique opportunity for *in silico* hypothesis generation via co-expression analysis. Here we leverage all published *V. cholerae* RNA-sequencing data, in combination with select data from other platforms, to generate a gene co-expression network that validates known gene interactions and identifies novel genetic partners across the entire *V. cholerae* genome. This network provides direct insights into genes influencing pathogenicity, metabolism, and transcriptional regulation, further clarifies results from previous sequencing experiments in *V. cholerae* (e.g. Tn-seq and ChIP-seq), and expands upon micro-array based findings in related gram-negative bacteria.

**Importance:** Cholera is a devastating illness that kills tens of thousands of people annually. *Vibrio cholerae*, the causative agent of cholera, is an important model organism to investigate both bacterial pathogenesis and the impact of horizontal gene transfer on the emergence and dissemination of new virulent strains. Despite this importance, roughly one third of *V. cholerae* genes are functionally un-annotated, leaving large gaps in our understanding of this microbe. Through co-expression network analysis of existing RNA-sequencing data, this work develops an approach to uncover novel gene-gene relationships and contextualize genes with no known function, which will advance our understanding of *V. cholerae* virulence and evolution.

## Introduction

Since the completion of the first *Vibrio cholerae* genome sequence in 2000, over a thousand *V. cholerae* isolates have been sequenced (1, 2). These sequences has allowed for the development of sophisticated phylogeographic models, which emphasize the importance of controlling the spread of virulent and antibiotic resistant *V. cholerae* strains to lower disease burden, in addition to fighting endemic local strains (2–6). The integration of hundreds of genomes paired with temporal and geographic information into ever growing phylogenies enables analyses using selection models to predict future population trends and derive biologically meaningful insights into *V. cholerae* evolution (7, 8). By developing treatment and vaccination strategies based on phylogenic models (9), organizations and governments can more efficiently leverage limited resources and more effectively prevent disease spread in line with the World Health Organization’s goal of eradicating cholera by 2030 (10).

Alongside advances in genomics research, the *V. cholerae* and broader bacterial biology communities have benefited greatly from other next generation sequencing (NGS) technologies. Targeted sequencing experiments have been essential in mapping complex virulence pathways, illuminating a novel interbacterial defense system, and expanding our knowledge of the role of non-coding RNA (ncRNA) in the vibrio life cycle (11–17). Further discoveries such as transcription factor mediated transposon insertion bias (18) and the role of cAMP receptor protein in host colonization (19) have benefited from composite research strategies utilizing multiple technologies. Similarly, meta-analyses utilizing pooled data from multiple experiments are empowered by the increasing availability of high quality bacterial NGS datasets. Expression data is particularly amenable to such pooling and can be used to accurately group genes into functional modules based on their co-expression (20). In bacteria, weighted gene co-expression network analysis (WGCNA) (21) has been successfully used to underscore biologically important genes and gene-gene relationships via “guilt-by-association” approaches (22, 23). These studies have taken advantage of larger and larger heterogeneous microarray datasets to provide novel biological insights via existing data.

Despite major advances in sequencing technologies and research strategies, most of the over two dozen existing RNA-seq experiments in *V. cholerae* have been limited to targeted approaches that involve quantifying the differential abundance of genetic material across a handful of conditions. Via these approaches, any change in expression observed in one experiment is nearly impossible to generalize to other treatment conditions and analyses are limited to a few pathways or genes of interest. In contrast, meta-analyses such as WGCNA can uncover much broader relationships throughout the entire genome by combining information from multiple datasets. As there is no existing co-expression analysis in *V. cholerae* to date, the accumulation of over 300 publicly available RNA-seq samples from targeted RNA-seq experiments represents a heretofore untapped resource for the cholera community.

Motivated by the success of pooled genetic sequencing analyses, our current work utilizes all publicly available *V. cholerae* RNA-seq based expression-level data to generate a co-expression network. We expand upon existing bacterial WGCNA approaches by integrating broader sequencing data (including ChIP-seq and Tn-seq) and multiple annotation platforms into our analysis. Our network ultimately contributes information on connections across all *V. cholerae* genes, including the roughly 1500 predicted but functionally un-annotated genetic elements that account for some 37% of the genome. More specifically, we implicate new loci in virulence regulation and clearly demonstrate a powerful and accurate approach to hypothesis generation via our described network.

## Results

### Gene network generation

To generate our co-expression analysis in *V. cholerae*, we applied our WGCNA pipeline to analyze twenty-seven *V. cholerae* RNA sequencing experiments deposited in NCBI’s Sequence Read Archive (SRA) in addition to two novel experiments. The RNA sequencing samples are derived from experiments exploring a range of important *V. cholerae* processes including intestinal colonization, quorum sensing, and stress response. In total, our network includes 300 individual RNA-seq samples (supplementary table S1). All samples were mapped to a recently inferred *V. cholerae* transcriptome derived from the N16961 reference genome (1, 13). This reference was chosen because the majority (293) of samples were collected from strains N16961 or the closely related C6706 and A1552.

Figure 1 outlines the process used to generate our co-expression network with a small subset of genes. Loci VC0384-VC0386 are known to be involved in cysteine metabolism while the two genomically adjacent loci VC0383 and VC0388 do not share this function. Following normalization of mapped transcripts (Fig. 1A), a weighted gene co-expression network analysis was performed using WGCNA (21). First, a Pearson correlation matrix is calculated for expression levels of all genes (Fig. 1B). This correlation matrix clearly captures strong relationships between co-expressing genes such as VC0384-VC0386 but can produce background noise from un-related gene pairs. We limit this noise by calculating a topological overlap matrix (TOM) (24) that weights pairwise co-expression data based on each gene’s interactions with all other genes (Fig. 1C). In this way, the relationships between genes that fall within the same subnetwork, i.e. VC0384-86, are favored while the signal from unrelated genes, i.e. VC0383 and VC0388, is abated. This TOM, after filtering for normalized values greater than 0.1, is used to construct an accurate co-expression network that captures biologically meaningful relationships (Fig. 1D).

**Figure 1:**
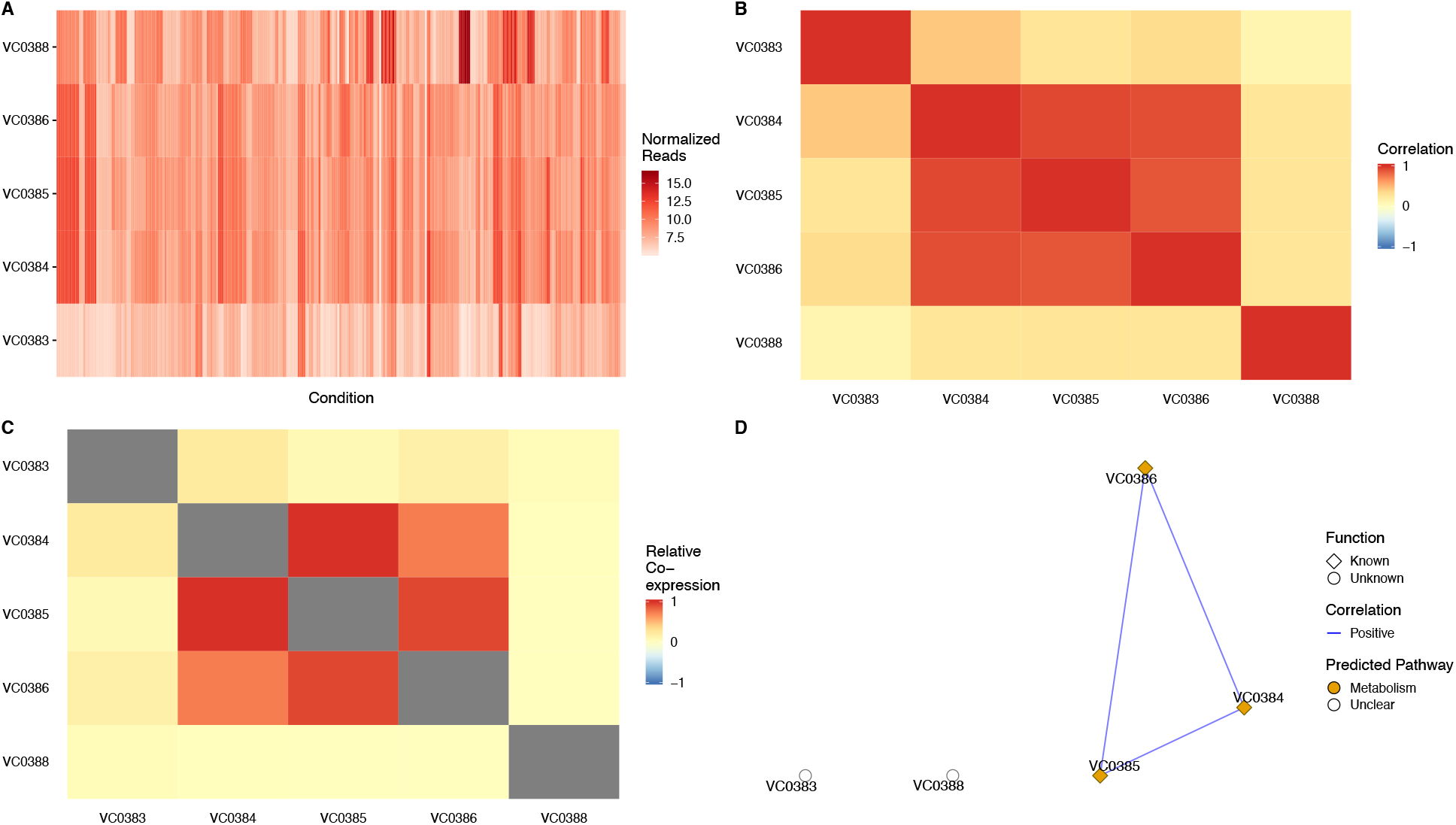
General outline of network construction. 1A) Normalized (log2) expression reads for the same genes across multiple conditions supply the basis for our co-expression analysis. In this small example, it is clear that genes VC0384-VC0386 have a very similar expression pattern across conditions. 1B) Correlations are calculated from the normalized counts in A for every pair of genes. The pattern seen in A becomes much clearer when looking at the correlation. 1C) An adjacency matrix (not shown) is calculated from the correlations in B and ultimately used to produce a topological overlap matrix (TOM) that supplies network edge weights with less noise than the raw correlation matrix. While the single of co-expressing pairs is dampened slightly, this step greatly decreases spurious relationships as it favors transcripts which coexpress with similar sets of genes rather than potentially noisy direct correlations. 1D) The final network groups transcripts that tightly co-express while indicating what pathway they are involved in. This network also includes functional and essentiality based knowledge. In this case, the three genes involved in cysteine metabolism (VC0383-VC0385, *cysHIJ*) form a subnetwork while the other genes do not meet our 0.10 co-expression cutoff.

In addition to co-expression data, our network and analyses incorporate information from multiple other sources. Our network includes predicted pathway annotations and gene functional knowledge from the NCBI Biosystems database as well as the DAVID, Panther, and KEGG databases (25–28). Additionally, importance labels are applied to genes with no known function which have been implicated as playing a role in intestinal colonization or *in vitro* growth via Tn-seq based essentiality experiments (14, 29). Information from ChIP-seq binding assays and microarray results were incorporated in downstream analyses to substantiate network derived relationships. By combining all of these data sources we were able to develop and analyze an informative network of co-expressing genes that provides both qualitative and quantitative information about relationships between transcripts across forty-nine gene-clusters covering the entire *V. cholerae* genome (Supp. Data S1-2).

### Genes in known pathways cluster together and contextualize genes of unknown function

As proof of the accuracy of our approach, we have highlighted four clusters that recapitulate known interactions between transcripts involved in specific pathways or cellular processes (Fig. 2). The correct grouping of transcripts encoding products such as ribosomal proteins, amino acid synthesis proteins, and tRNA transcripts that have largely known functions and are involved in well-studied, highly conserved cellular processes provides a positive control for the validity of our network clusters (Fig. 2A-C). Likewise, the clustering of genes known to be involved in more specialized processes such as biofilm formation (Fig. 2D) further underscores the success of our approach.

**Figure 2:**
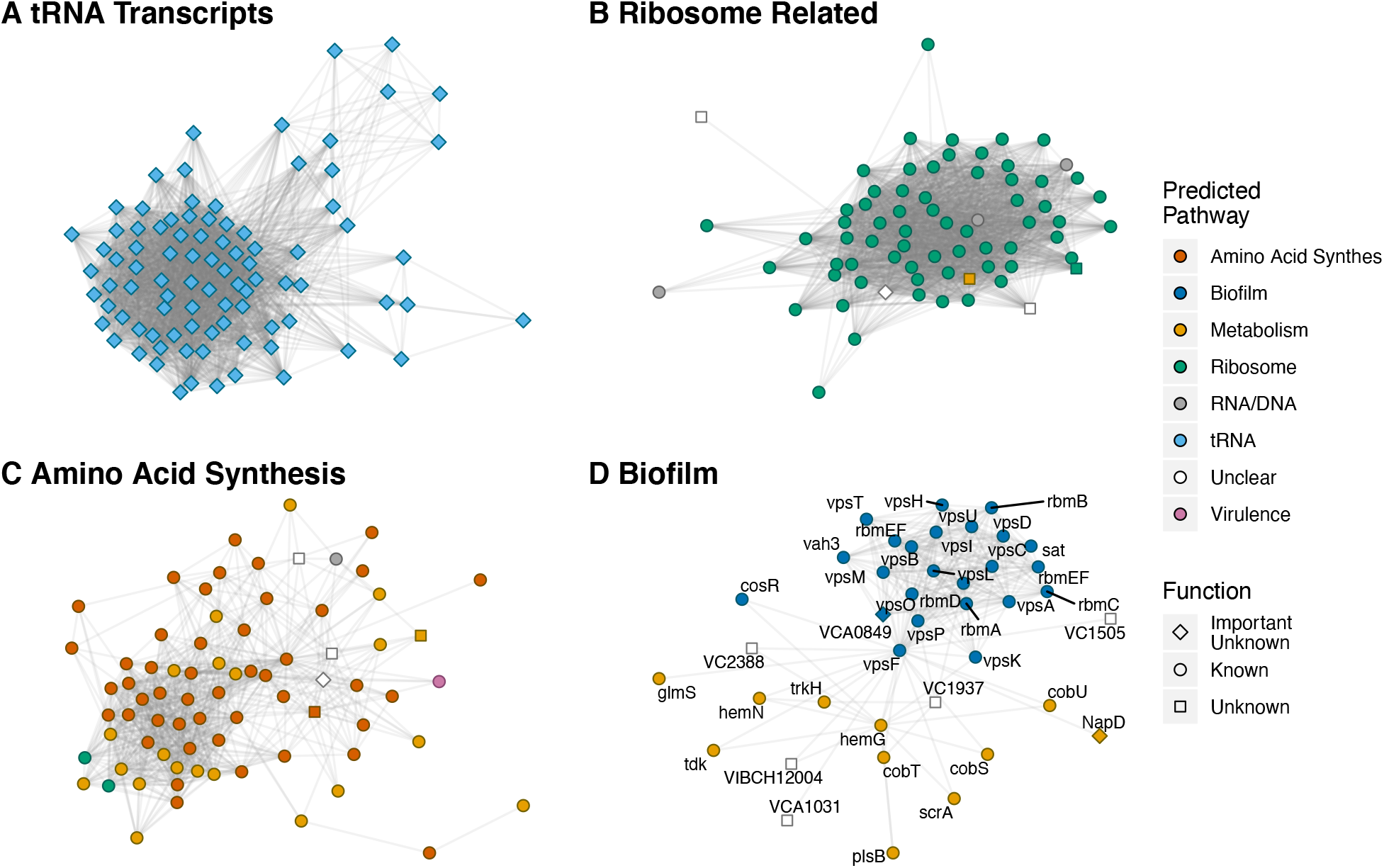
Sub-networks recapitulating known results The four depicted subnetworks each contain subsets of transcripts which are known to be largely involved in the same biological process. For each subnetwork, the nodes represent transcripts while the edges represent a co-expression relationship of at least 0.1 between transcripts. A) This sub-network consists completely of tRNA transcripts. B) These transcripts are almost completely related to ribosomal structure and/or function. C) These transcripts play a role in amino acid synthesis. D) This sub-network contains a majority of transcripts that play a role in biofilm formation in addition to unrelated genes.

The subnetworks mentioned above also support the utility of our analysis in powering guilt-by-association based inference of gene function (30). Because each of these gene clusters contain co-expressing genes that are involved in the same biological process, it can be assumed that unannotated genes in the same cluster are likely involved in the same process. Such links, while not definitive on their own, can be used with other data to hint at gene functions. For example, genes with known function in Fig. 2D are primarily involved in biofilm formation (31, 32). This clustering of biofilm genes suggests that the few genes with no known function in this subnetwork may be involved in the same process. Two of these unannotated transcripts, VC1937 and VC2388, are, per GO cellular component location labels, “integral membrane components.” Further, the VC2388 locus is directly upstream of a Vcr084, a short RNA involved in quorum sensing which is essential for biofilm formation (33). Taken together, this evidence suggests that VC1937 and VC2388 may play a role in some of the complex membrane restructuring necessary for biofilm formation. In facilitating such guilt-by-association approaches to novel hypothesis generation, our co-expression network serves as a highly efficient substitute for more traditional screening assays.

### A virulence subnetwork suggests novel gene functions

While the biofilm associated subnetwork presents a relatively simple example of the functional insights our co-expression data can yield, the virulence-related subnetwork (Fig. 3A) represents a more complex case in which genes of known function provide clues to the role of unannotated genes. The majority of transcripts in this module originate from within the virulence-related ToxR regulon that consists principally of genes on the *V. cholerae* pathogenicity island 1 (VC0809-VC0848) and cholera toxin sub-units A and B (*ctxAB*, VC1456 and VC1457) (34). Other genes in this subnetwork, such as *vpsJ*, VC1806, VC1810, and chitinase, are predominately localized to virulence islands and other areas of the genome under tight control of the known virulence regulators ToxR, ToxT, or H-NS as determined via ChIP and/or RNA-seq (35–37). The clustering of such genes with well-characterized interactions into a cohesive subnetwork is further validation of our ability to generate accurate co-expression maps of related genes. The association of uncharacterized genes in these clusters suggests they may also play a role in *V. cholerae* virulence and generates hypotheses about the function of unknown genes within this module.

**Figure 3:**
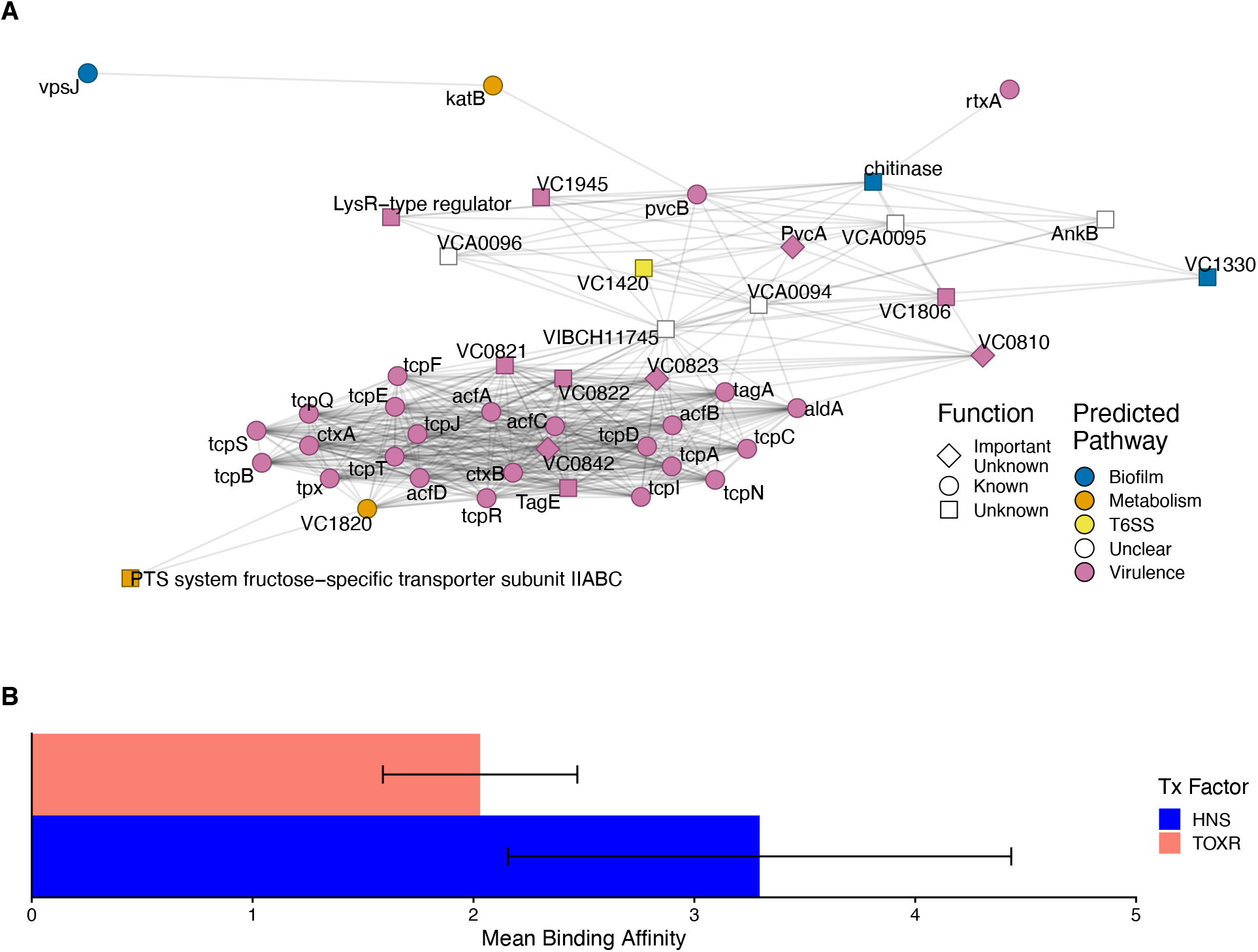
Virulence related subnetwork. 3A) This subnetwork contains a majority of genes that are predicted to be involved in virulence related pathways, providing clues to the genes with no known functions such as those at locus VCA0094-VCA0096. 3B) Mean binding affinity (log2 fold change in occupancy compared to loading control) for different virulence-associated transcription factors near the VCA0094-96 locus. Both HNS and TOXR show a significant binding preference for this region. Error bars indicate standard deviation from the mean.

Many of the important transcripts with unknown function are expected to co-express with known virulence genes because they fall within vibrio pathogenicity island (VPI)-1 (VC0810, VC0821-VC0823, VC0842) or VPI-2 (VC1806, VC1810), or are proximal to other virulence genes (VC1945) (38, 39). However, our analysis also identified genes such as VCA0094-VCA0096 which are on a completely different chromosome than the rest of the subnetwork and do not neighbor any known virulence elements.

A major benefit of our approach is that we incorporate additional regulatory data such as ChIP and Tn-seq into our co-expression analysis, allowing us to verify the association between VCA0094-VCA0096 and virulence pathways using existing experimental data. Tn-seq analysis has previously identified VCA0094 and VCA0095 as essential for infection of a rabbit intestine (14), suggesting that these loci play a role in virulence. Because transcripts for these genes co-express with genes regulated by ToxT, ToxR, and H-NS, we also probed existing ChIPseq binding datasets (12, 19, 35) to see if any of these well-studied transcription factors bind near the VCA0094-96 loci. While ToxT binding was not observed near this site (data not shown), our analysis identified significant peaks in the promoter region of VCA0094 for both ToxR and H-NS as calculated via re-analysis of existing binding data from (35). Both peaks showed a large and significant increase in binding affinity (log_2_ fold change in average occupancy) when compared against input controls (Fig. 3B). H-NS showed a clear binding peak in the region of the VCA0094 promoter that extended in a diffuse manner to the VCA0095 TSS while ToxR binding covered a similar region but was more diffuse throughout (data not shown). Collectively these results indicate virulence related functions for the products of the VCA0094-VCA0096 transcripts. Although the exact mechanistic role of these genes remains elusive, we have nevertheless demonstrated the ability of our pipeline to generate meaningful hypotheses by incorporating existing data from a multitude of sources.

### Co-expression data provides an accurate *in silico* complement to RNA-seq

In addition to the guilt-by-association inference described above, co-expression analysis can provide a partial substitute or complement to RNA-seq experiments. Novel, meaningful genetic relationships can be found in a co-expression network by focusing on the transcripts that are co-regulated with a gene of interest.

We can apply a network-based approach in lieu of new RNA-seq based experiments to identify genes which co-express with *rpoS* (VC0534) and are similarly involved in bacterial stress response. As our network utilizes only RNA-seq based transcriptomics studies and none of these studies involves direct manipulation of *rpoS*, we can compare existing microarray data involving an *rpoS* (VC0534) deletion mutant (40) to determine how accurate our approach is. When applying an absolute co-expression cutoff of 0.1, 273 genes are identified as having a relationship with *rpoS* expression in both our network analysis and the *rpoS* mutant microarray data (Fig. 4A). This represents nearly two-thirds of genes identified as differentially expressed in the original microarray study. While our network links far more genes with *rpoS* than the microarray approach, this is in line with recent RNA-seq based work that found that 23% of the E. coli genome is regulated by RpoS (41). Additionally, all of the flagella and chemotaxis related proteins highlighted as particularly informative in the original study are identified by our analysis (Fig. 4B) and relevant values (i.e. network co-expression and microarray-derived log fold change in expression) for the 273 shared transcripts have a Spearman correlation of −0.516. This accuracy is achieved without any direct genetic manipulation of the *rpoS* locus in the RNA-seq datasets used to generate our co-expression network and serves as a testament to the potential utility and versatility of our approach.

**Figure 4:**
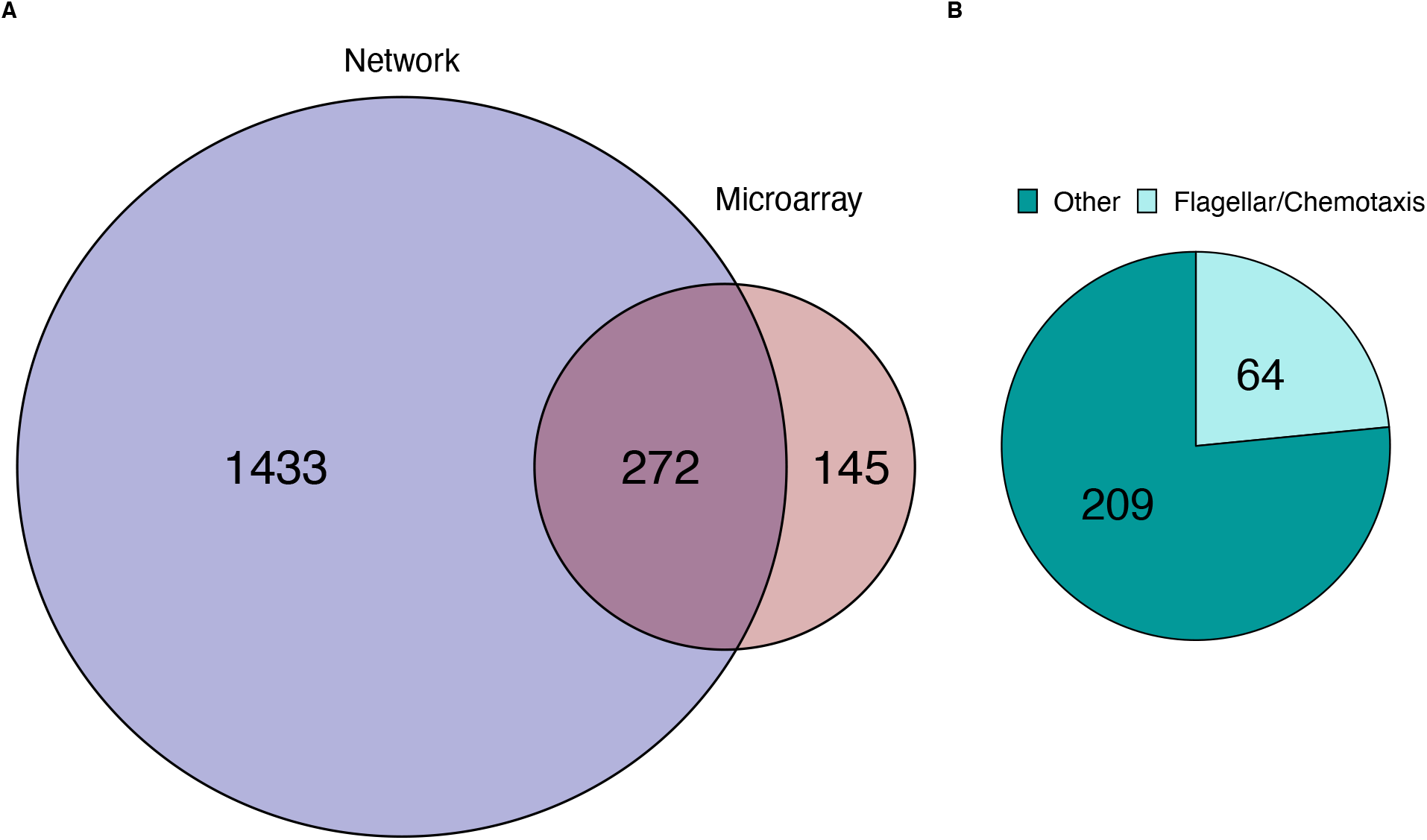
Comparing RpoS microarray data to co-expressing genes in our WGCNA A) Overlap of genes with expression pattern related to *rpoS* expression as identified via our network analysis (blue) and existing microarray data (red). The overlapping region identifies 272 genes that are common between the two analyses. B) Breakdown of shared genes (overlapping region in A). All of the flagellar and chemotaxis genes highlighted as particularly important in the microarray dataset are identified by both methods.

Our approach to isolating genetic interactions also has advantages over transcriptomics-focused sequencing. As seen in Fig. 4A, our network-based analysis identifies far more genes associated with *rpoS*. This is likely because RNAseq-based approaches are can identify a broader range of gene transcripts as they are not limited by restrictive microarray probes (42). Separate from differences in underlying technology, co-expression networks are also more likely to detect genes regulating a target’s expression than traditional transcriptomics experiments which largely capture downstream responses to changes in a target’s expression (43, 44). Thus, a co-expression network can provide an alternative perspective to complement or clarify transcriptomics data.

## Discussion

We have successfully constructed the first *V. cholerae* co-expression network through a computationally inexpensive process that is simple, easily expanded upon, and straightforward to implement in other organisms. Our network effectively identifies canonical gene clusters related to specific molecular pathways or functions, such as those corresponding to rRNAs or biofilm proteins. We have also outlined two use-cases for the data provided and have shown the accuracy of both approaches either experimentally or using existing data. Additionally, we have included relevant network files as well as raw read counts across RNA-seq conditions (Supp. Data S1-2 & Supp. Table S2) alongside all code used in our analysis (see Materials and Methods) to encourage broad usage of this data.

Our results have proven both the utility and accuracy of our approach despite in-depth analysis limited to a handful of genes across five of the forty-nine observed gene clusters. Furthermore, our work with the virulence subnetwork supports previously published research loosely implicating genes VCA0094-96 in virulence and virulence related functions. All three transcripts have shown up in screens focusing on biofilm development (45), and SOS response (13). From a mechanistic perspective, protein homology analysis via NCBI’s Conserved Domain Database (46) indicates that VCA0094 possesses a DNA-binding transcriptional regulator domain while VCA0096 contains domains that implicate it in protein activation via proteolysis. These data combined with our novel findings hint at the potential biological importance of this genomic locus.

When viewed through the lens of a specific gene of interest, co-expression data is in large part analogous to the differential expression data produced by RNA-seq experiments. While RNA-seq offers finer assay control and can be tailored more exactly to suit a specific research question, there are both technical and practical limitations that may make such an approach impractical. Whether an experimenter is interested in examining the role of an essential locus or is limited by available resources, our co-expression analysis presents a fast, free, and faithful alternative for probing genetic interactions as outlined in our analysis of *rpoS* above.

Major motivations for this work include the successful implementation of bacterial-focus, microarray-based co-expression networks and the lack of clear functional knowledge for a large portion of *V. cholerae* genes. Besides more simple guilt-by-association studies (22, 23), co-expression networks have helped to elucidate relationships in diverse microbial communities (47–50) and enable comparisons across strains and species (51–53). These works as well as the relative dearth of knowledge about the *V. cholerae* genome (roughly two third of genes are annotated compared to around 86% percent of all *E. coli* genes (54)) and the growing abundance of *V. cholerae* focused NGS data served as the impetus for this research.

The calculated co-expression network, though accurate, could be improved via the inclusion of more experiments and more extensive SRA annotations. Our somewhat limited pooled dataset consisting of three hundred samples is an order of magnitude off from the few thousand samples necessary to derive the most faithful co-expression estimates (55). Though sample size will improve as more *V. cholerae* RNA-seq experiments are published, more samples may also increase the risk posed by batch effects which cause spurious correlations among genes through technical variation (56, 57). The diverse structure of our current data helps to minimize the impact of batch effects but this would be offset by the future inclusion of larger datasets from single experiments. While automated sample clustering methods (58–60) can effectively group overly correlated samples, there is no way to know if the correlation is biological (i.e. meaningful) or technical (i.e. noise) in origin. Likewise, manual curation of batch annotations is also difficult since few SRA records are extensively annotated with detailed experimental conditions (e.g. bacterial growth stage, exact medium used). Thus, careful consideration may be necessary when expanding and generalizing this analysis to include future data.

The mapping of raw reads to a transcriptome derived from a single reference genome presents a limitation to our current work. While this approach is reasonable given the similarity of the vast majority of included strains to our reference, a more elaborate comparative transcriptomic strategy (61, 62) would be ideal if more diverse samples are included in future analyses. This is especially true when considering the inclusion of expression data from clinical samples which are likely to have much more genomic variability than the closely related lab cultured strains used to construct our network. On the other hand, because comparative transcriptomics requires defining homologous alleles across all strains analyzed (63), such an approach would greatly increase the difficulty of incorporating strains without an assembled genome.

In summary, our co-expression network can drive functional hypotheses for unannotated genes in *V. cholerae*. As the Vibrio community steadily adds high quality data from increasingly sophisticated sequencing experiments to public databases our imputed network can only improve, providing ever deeper insights into the *V. cholerae* genome. At the same time, highly annotated transcript-based co-expression networks can empower research with related technologies (e.g. single cell transcriptomics and dual RNA-seq) and research into a host of other clinically relevant bacteria, such as *Pseudomonas aeruginosa* or *Staphylococcus aureus* which have over 2000 and 1400 RNA-seq experiments in SRA respectively.

## Materials and Methods

### Data Collection and Processing

All RNA and ChIP sequencing data were downloaded from the Sequence Read Archive (SRA)(64) and converted to compressed fastq files using the SRA toolkit (https://www.ncbi.nlm.nih.gov/sra/docs/toolkitsoft/) (see Table S1 for details on included experiments). RNA-seq samples were selected by searching the SRA on Sept 10^th^, 2019 for the Organism and Strategy terms “vibrio cholerae” and “rna seq” respectively, resulting in 326 initial samples including the 34 novel samples from this publication (PRJNA601792). Samples were mapped to a recently inferred *V. cholerae* transcriptome derived from the N16961 reference genome (1, 13) using Kallisto version 0.45.1 (65). This reference was chosen because the majority (293) of samples were collected from strains N16961 or the closely related C6706 and A1552. 26 low quality samples with < 50% of reads mapping to the reference transcriptome were discarded before further analysis, leaving 300 samples used for further analysis.

For ChIP-seq analysis, accession numbers were identified via the relevant publications (12, 19, 35) and sequences were downloaded from SRA and converted to fastq files as above. Raw reads were mapped to the same N16961 reference genome using Bowtie 2 version 2.3.5.1 (66). From this mapping, peaks were identified using MACS2 version 2.1.2 with an extsize of 225 (various sizes from 150 to 500 were tested with little observable difference in peaks identified) (67) and differential binding and significance were calculated using DiffBind version 2.12.0 (68).

Processed Tn-seq data were collected directly from published datasets. *In vitro* essentiality and semi-essentiality labels were derived from Chao et al. 2013 Table S1 (29), with the original labels of domain essential and sick genes replaced with essential and semi-essential respectively. We used Table S2 from Fu, Waldor, and Mekalanos 2013 (14) to label genes involved in host infection, with any gene exhibiting a log_2_ fold change less than negative three deemed essential and any gene with a log_2_ fold change between negative one and negative three deemed semi-essential.

### Network Construction

Figure 1 highlights the process used to generate our co-expression network. Kallisto derived reads were first imported into R via tximport (69), then normalized using DESeq2 version 1.24.0 (70), resulting in values that are comparable across conditions and experiments. Following normalization, a weighted gene co-expression network analysis was performed using WGCNA (21). This process is highlighted with a subset of data in Figure 1 and consists of the sequential calculation of a Pearson correlation matrix, adjacency matrix with power ß=6, and, ultimately, topological overlap matrix (TOM) (24) from normalized gene expression counts across conditions. We further filtered this TOM to exclude samples with weighted co-expression <0.1 for all analysis included in the Results section.

Predicted pathway annotations and gene functional knowledge are derived from the NCBI Biosystems database as well as DAVID, Panther, and KEGG databases (25–28). Genes lacking functional knowledge which are identified as essential or semi-essential in either Tn-seq dataset are labeled in network visualizations as “important.”

### Data Availability

SRA accession numbers and information on included samples can be found in Supplementary Table S1. A full, unfiltered network graph is provided in Supplementary File S1 with the corresponding node labels in Supplementary File S2. Raw, un-normalized read counts are also provided in Supplementary Table S2 All data analysis and figure generation were done using the R programming language, with code available at DOI: 10.5281/zenodo.3572870.

